## Supplementary Figures for "Full-length direct RNA sequencing uncovers stress-granule dependent RNA decay upon cellular stress"

### Supplementary Figure 1

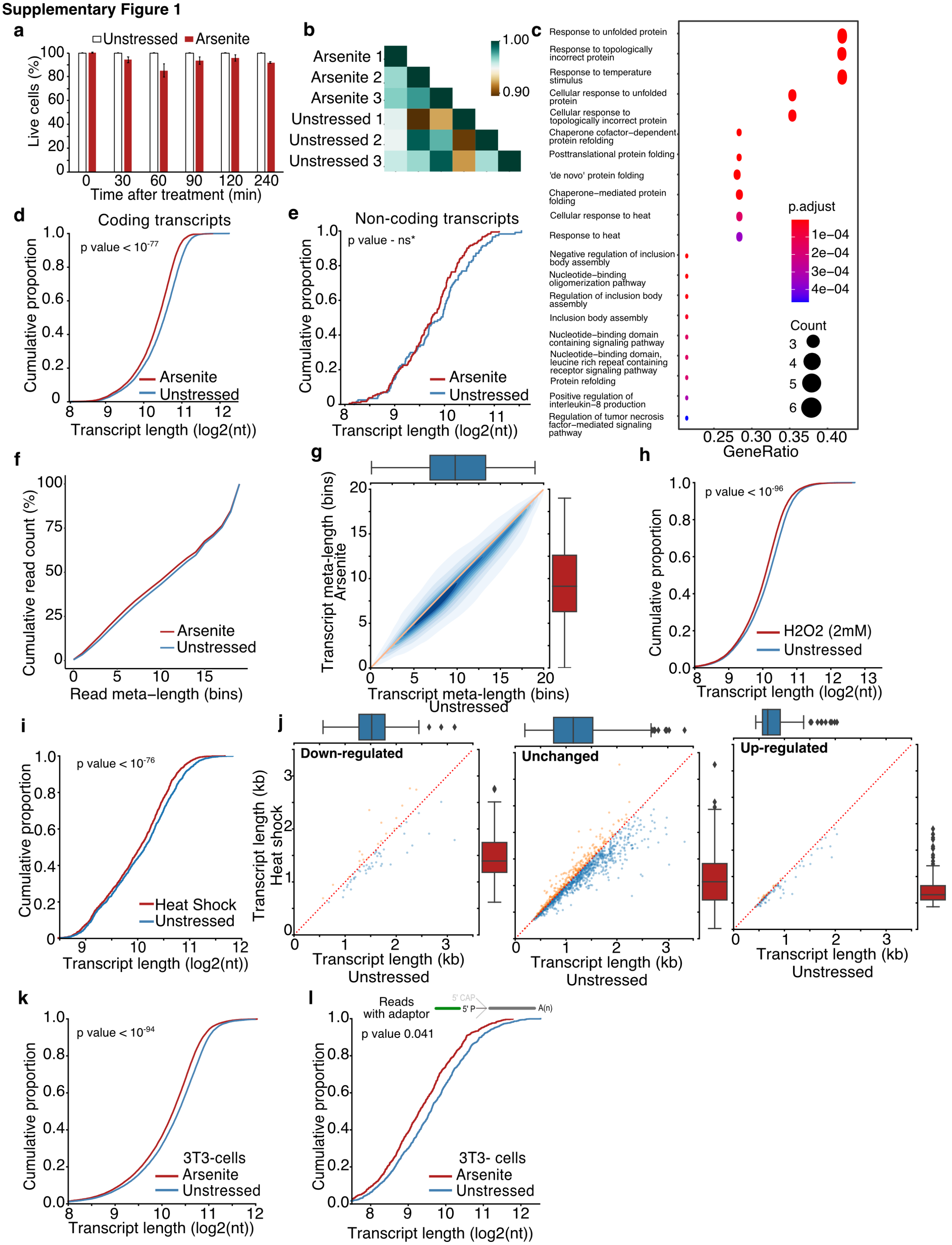

**Sup. Fig 1: RNA shortening upon cellular stress.** **a)** MTS cell proliferation assay showing the percentage of live cells (y-axis) upon arsenite treatment compared to unstressed at different time points (x-axis). Error bars represent std for biological replicates (n = 2). **b)** Correlogram of transcript expression for unstressed and arsenite-treated cells. **c)** Gene ontology for biological processes for differentially expressed genes in arsenite-treated versus unstressed cells. **d-e)** Cumulative distribution plot of average transcript length for coding (d) and non-coding (e) transcripts. Transcripts with less than 5 reads have been removed. **f)** Cumulative distribution of read meta-length for arsenite-treated and unstressed cells. **g)** Contour density plot of average transcript meta-length for arsenite-treated and unstressed cells. Only reads with an identified poly(A) tail are used. **h)** Cumulative distribution plot of average transcript length for H2O2-treated cells and unstressed. **i)** Cumulative distribution plot of average transcript length for heat shock cells and unstressed. **j)** Scatter plots of average transcript length for heat shock against unstressed cells stratified by differential expression change. Down-regulated: (-Inf, -0.5), Unchanged: (-0.5, 0.5), Up-regulated (0.5, Inf) fold-change. Only transcripts with at least 5 aligned reads are shown. Red dotted indicates the y=x line. Color indicates transcripts below (blue) and above (orange) the diagonal. **k-l**) Cumulative distribution plot of average transcript length for the mouse 3T3 cells treated with arsenite and unstressed. All reads (k) or only reads with ligated adaptor (l) are used.

**Supplementary Figure 2**

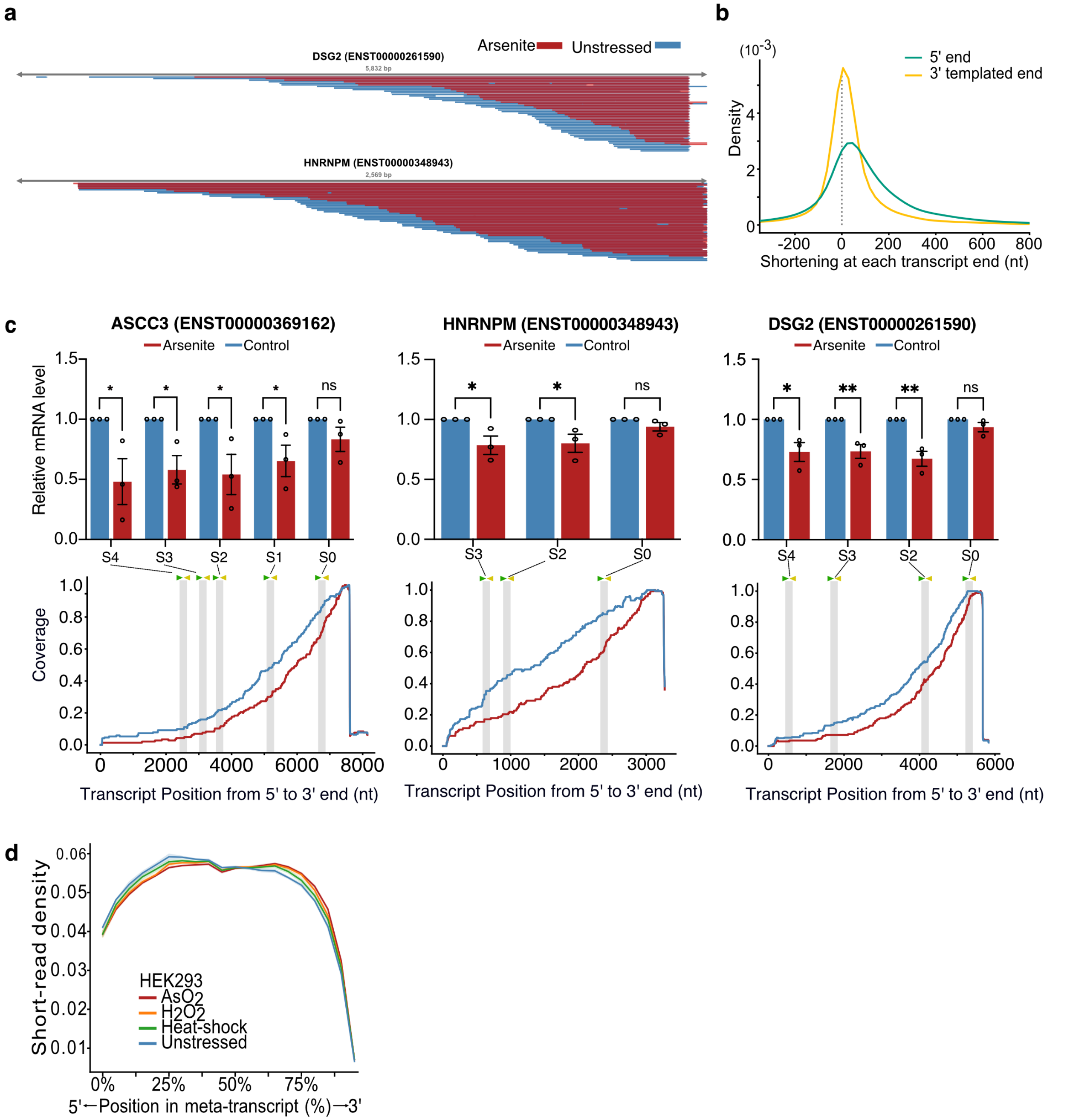


**Sup. Fig. 2: Characterization of stress-induced shortened RNAs identified by NanopLen. a)** IGV screenshot of aligned reads in arsenite-treated (red) and unstressed cells (blue) for significantly shortened transcripts *HNRNPM* and *DSG2*. Libraries were randomly downsampled to maximum 50 reads per window and the libraries were overlayed. All reads were used, irrespective of adaptor ligation status. **b)** Density plot of transcript shortening at the 5′ and 3′ templated end. Positive values on X-axis indicate that the 5′ or 3′ of transcripts in arsenite-treated cells are correspondingly downstream or upstream of their unstressed counterparts. **c)** RT-qPCR (top) and read coverage (bottom) for transcripts ENST00000369162 of gene ASCC3, ENST00000261590 of gene DSG2 and ENST00000348943 of gene HNRNPM for arsenite-treated and control cells. PCR primer pairs were designed for varying locations along the transcript and their approximate location is marked on each plot. Signals were normalized against ACTB. Error bars correspond to SEM (N = 3) and individual points are shown. P-values were calculated using t-test. **: P <0.01and *:P<0.05. **d)** Meta-plot of short-read RNA-Seq density on all transcripts for unstressed, $AsO_2$, $H_2O_2$ and heat shock -treated cells. Shade indicates standard error of the mean for replicates.

**Supplementary Figure 3**

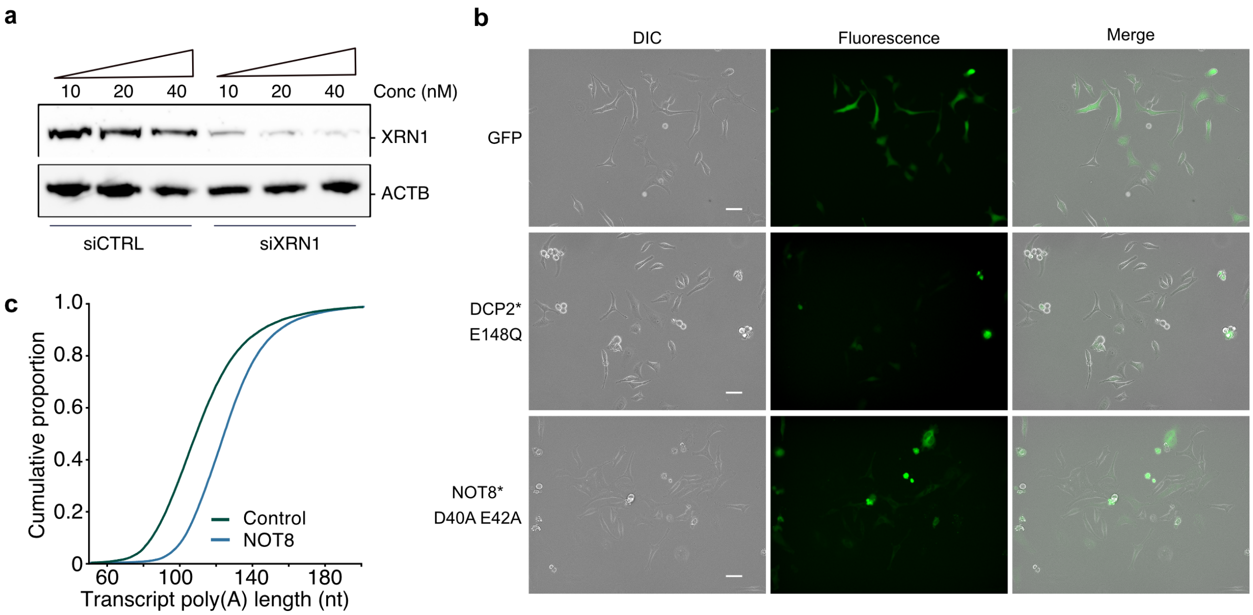


**Sup. Fig. 3. The effect of XRN1 knockdown on RNA shortening. a)** Immunoblot for XRN1 for cells transfected with mock and *XRN1*-targeting siRNA at varying concentrations. ACTB is used as control. **b)** Visualization by epifluorescence of GFP, GFP-fused DCP2* E148Q and GFP-fused NOT8* D40A E42A expression in HeLa cells. Scale bar 50 µm. **c)** Cumulative distribution of average transcript poly(A) tail length for NOT8*-expressing and GFP control cells.

**Supplementary Figure 4**

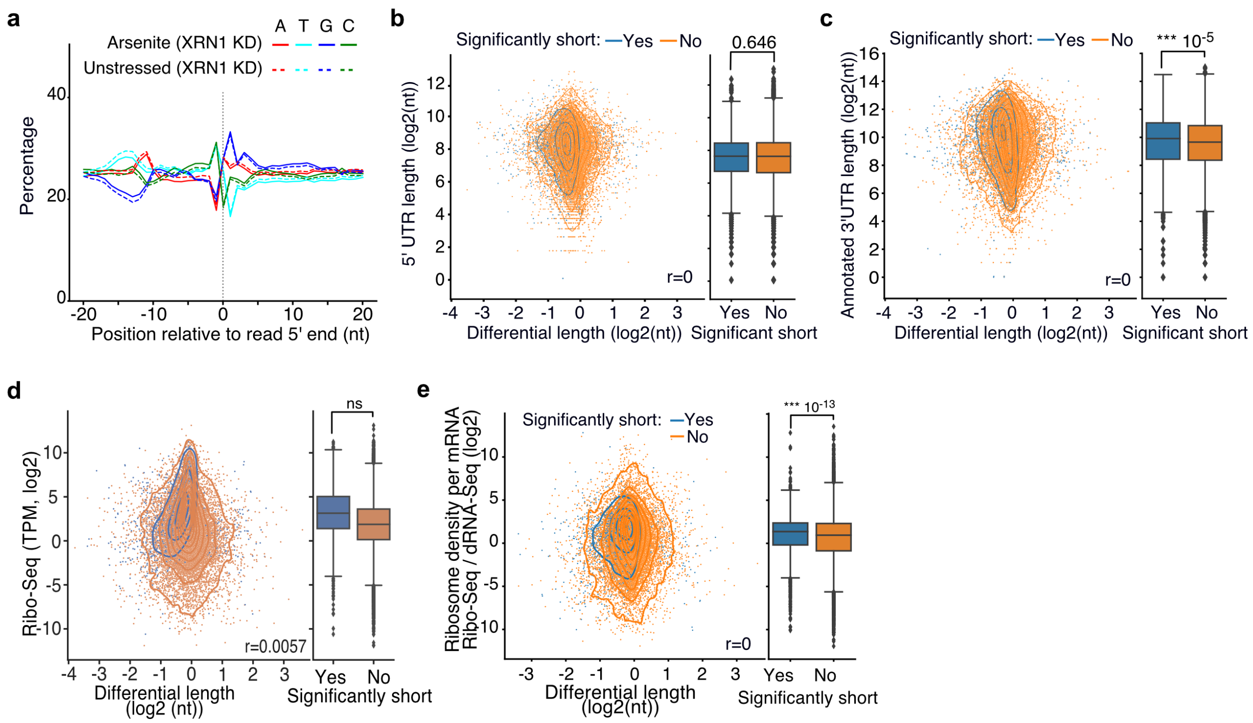


**Sup. Fig. 4: Association of cis-regulatory elements and ribosome occupancy with RNA shortening. a)** Nucleotide composition around the 5′ end of reads in arsenite-treated and unstressed cells upon XRN1 silencing. All reads were used, irrespective of adaptor ligation status. **b-c)** Scatter plot of annotated 5′ (a) and 3′ (b) UTR length against transcript differential length in arsenite-treated and unstressed cells. The box plots on the right side summarize the y-axis variable for significant and non-significantly shortened transcripts. **d-e)** Scatter plot of ribosome profiling footprint (d) and translational efficiency (e) levels against differential length in arsenite-treated and unstressed cells. Transcripts are stratified by significance of shortening. The Pearson’s correlation coefficient and the Mann-Whitney-U test p-value are shown.

**Supplementary Figure 5**

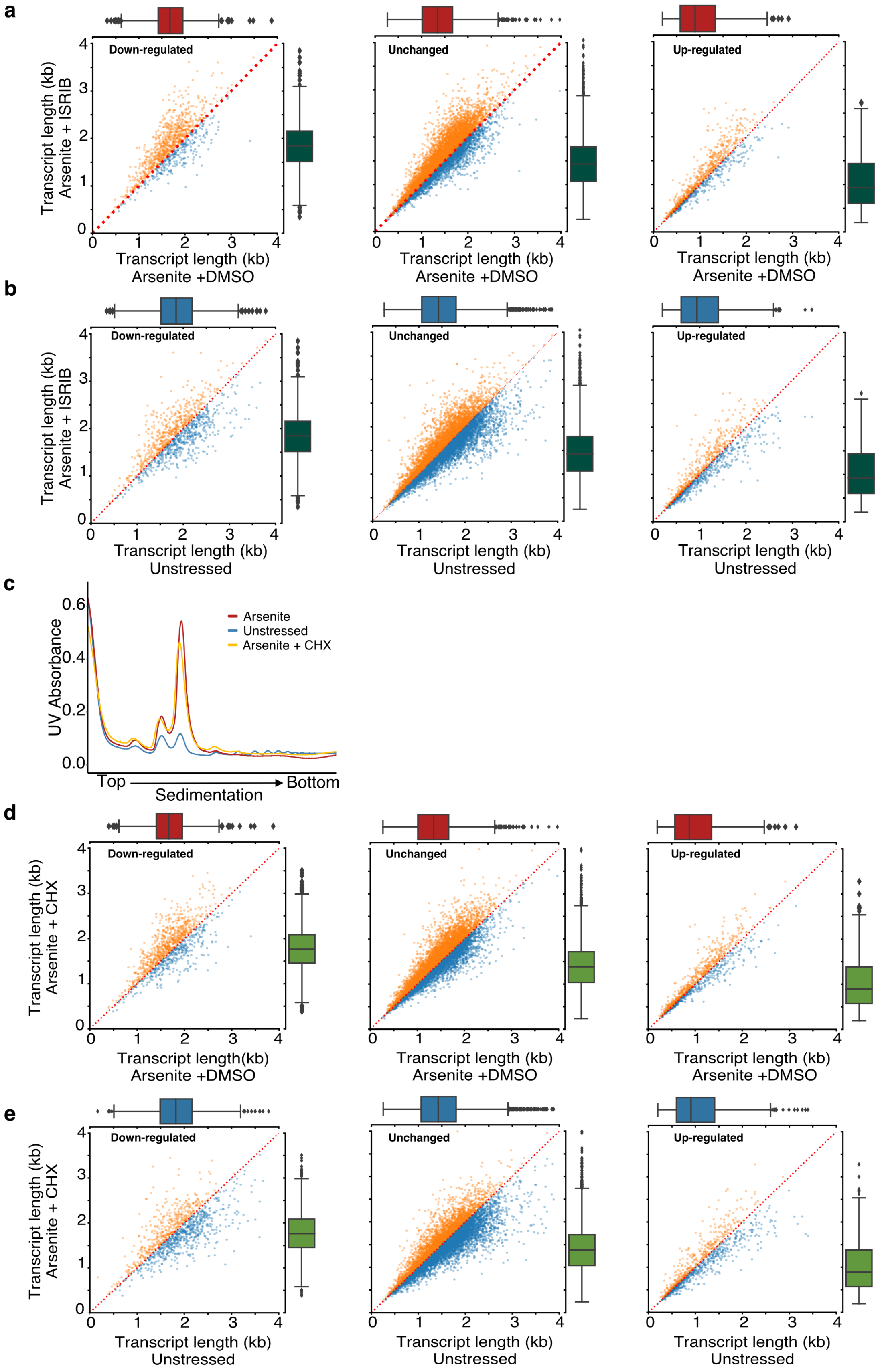

**Sup. Fig. 5. Translation and RNA shortening. a)** Scatter plots of average transcript length for arsenite-treated cells with and without ISRIB stratified by their differential expression change. Down-regulated: (-Inf, -0.5), Unchanged: (-0.5, 0.5), Up-regulated (0.5, Inf) fold-change. Only transcripts with at least 5 aligned reads are used. Red dotted indicates the y=x line. Color indicates transcripts below (blue) and above (orange) the diagonal. **h)** Same as (g) for arsenite and ISRIB treated cells against control. **c)** Ribosome sedimentation curve following cell treatment with cycloheximide (CHX) (25 μg/ml) for arsenite-treated and unstressed cells. **d-e)** Same as (a) and (b) for cycloheximide CHX instead of ISRIB.

**Supplementary Figure 6**

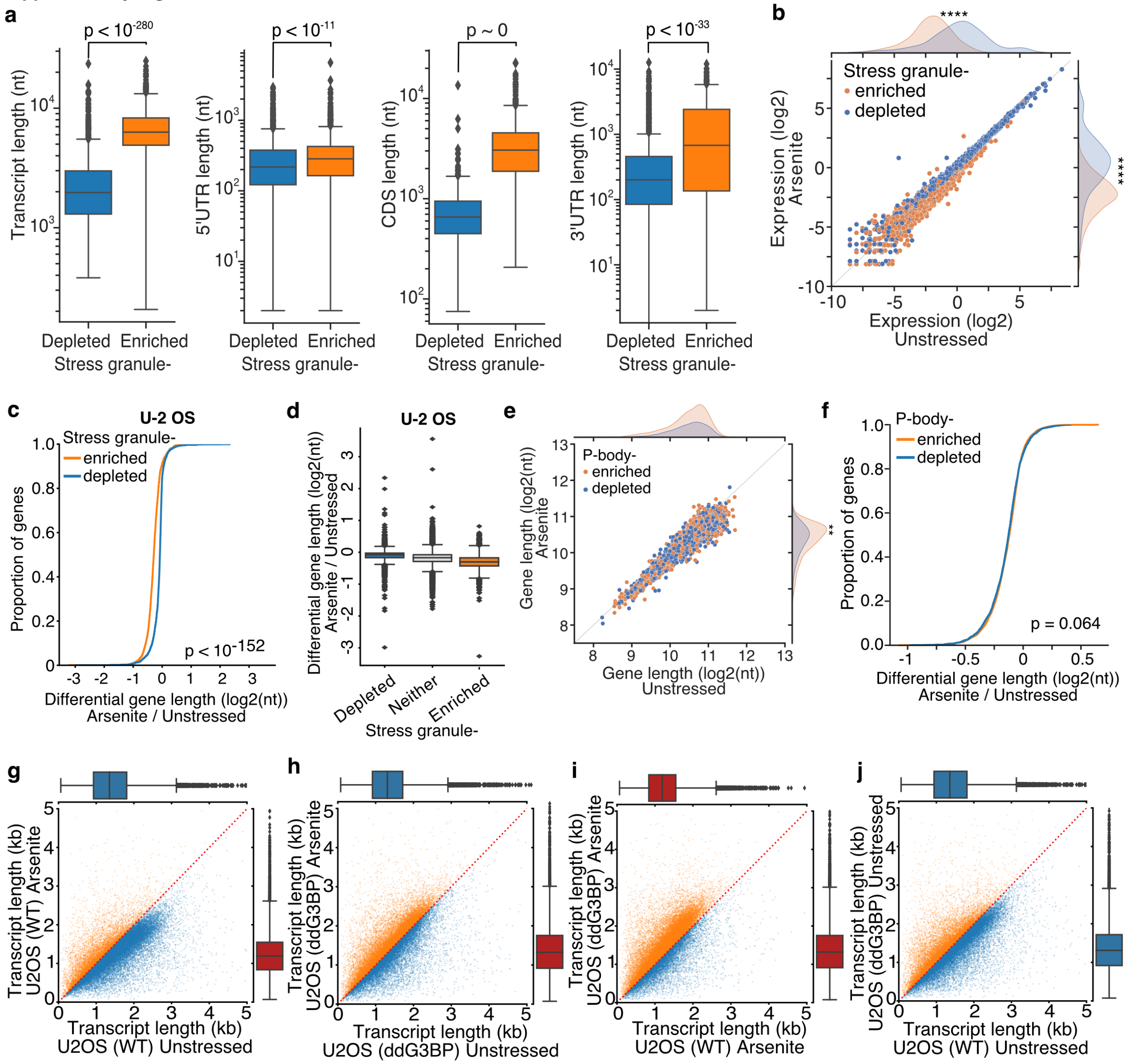


**Sup. Fig 6: Inhibition of stress-granule formation in cells devoid of G3BP1/2 rescues RNA decay. a)** Box plots of annotated total, 5′ UTR, coding sequence and 3′ UTR length for SG enriched and depleted transcripts. **b)** Scatter plot of upper 90th quantile normalized gene expression for arsenite-treated and unstressed cells stratified by gene SG localization. **** indicates p-value < $10^{-172}$. **c)** Cumulative density plot of transcript differential length for SG enriched and depleted gene transcripts in U-2 OS cells. Only transcripts of coding genes are used. **d)** Box plots of differential gene length in arsenite-treated and unstressed cells stratified by SG enrichment in U-2 OS cells. **e)** Scatter plot of average gene length for arsenite-treated and unstressed cells stratified by gene P-bodies localization. ** indicates p-value < 0.01. **f)** Cumulative distribution plot of average gene length difference in arsenite-treated and unstressed cells stratified by gene P-bodies localization. **g-j)** Scatter plot of average transcript length for arsenite-treated and unstressed U-2 OS and ΔΔG3BP1/2 cells. Only transcripts with at least 5 aligned reads are used. Red dotted indicates the y=x line. Color indicates transcripts below (blue) and above (orange) the diagonal.
